# Individual repeatability and heritability of divorce in a wild population

**DOI:** 10.1101/254151

**Authors:** Ryan R. Germain, Matthew E. Wolak, Jane M. Reid

**Affiliations:** School of Biological Sciences, University of Aberdeen, Aberdeen, UK

**Keywords:** indirect genetic effects, mating system evolution, social monogamy, quantitative genetics

## Abstract

Understanding micro-evolutionary responses of mating systems to contemporary selection requires estimating sex-specific additive genetic variances and cross-sex genetic covariances in key reproductive strategy traits. One key trait comprises the occurrence of divorce versus mate-fidelity across sequential reproductive attempts. If divorce represents an evolving behavioural strategy it must have non-zero individual repeatability and heritability, but quantitative estimates from wild populations are scarce. We used 39 years of individual breeding records and pedigree data from free-living song sparrows (*Melospiza melodia*) to quantify sex-specific permanent individual and additive genetic variances, and hence estimate repeatability and heritability, in liability for divorce. We estimated moderate repeatability in females, but little repeatability in males. Estimates of additive genetic variance were small in both sexes, and the cross-sex genetic covariance was close to zero. However, the total heritability was likely non-zero but small, indicating low potential for micro-evolutionary response to selection. Rapid micro-evolution of divorce rate therefore appears unlikely, even if there were substantial fitness benefits of divorce and resulting selection.

## INTRODUCTION

Pair-bond resilience, resulting from mate-fidelity (i.e. maintaining pair-bonds over multiple breeding attempts) versus divorce (i.e. dissolving pair-bonds to re-pair with a new mate) [1,2], is a key feature of animal mating systems that results from sexual selection and can fundamentally influence the distribution of offspring across parents and the resulting population-wide distribution of fitness. Divorce occurs in numerous socially-monogamous taxa and may increase individual fitness by counter-acting constraints on initial mate choice, and hence be adaptive [2,3]. However, if there is to be ongoing adaptive micro-evolution of divorce rate in response to selection [2,4–6] then individual liability for divorce must have non-zero heritability, and hence vary consistently among individuals (i.e. have non-zero individual repeatability).

In general, repeatable expression of mating and reproductive traits implies that selection may act consistently on individuals over their lifetimes, and defines the maximum possible heritability [7,8]. Decomposing total phenotypic variance into repeatable individual variance (V_I_), additive genetic variance (V_A_) and environmental variance allows estimation of heritability and, in principle, indicates the potential for micro-evolutionary responses to contemporary selection [9]. However, divorce represents an interesting class of jointly-expressed (i.e. ‘emergent’) traits that result from female-male interactions, and hence can be simultaneously influenced by genetic effects of both sexes. Further, correlated expression of sex-specific genetic effects, leading to non-zero cross-sex genetic covariance (COV_A♀♂_), can generate evolutionary sexual conflict [10] and further alter the potential for evolutionary responses to selection on either or both sexes that could result from sex-specific fitness consequences of key mating decisions (e.g. [11]). Understanding the evolutionary dynamics of divorce, and the implications for other co-evolving reproductive behaviours contributing to mating systems, therefore requires explicit estimation of sex-specific V_A_ (V_A♀_ and V_A♂_), and COV_A♀♂_, in divorce in populations experiencing un-manipulated natural and sexual selection environments.

Advances in quantitative genetic methods mean that V_A♀_, V_A♂_, and COV_A♀♂_ underlying emergent traits can be estimated given complex relatedness structures arising in wild populations (e.g.[9]). Since divorce versus mate-fidelity represent alternative outcomes of pairing decisions across consecutive breeding attempts, divorce is appropriately modelled as a ‘threshold trait’, where breeding pairs’ underlying continuous joint liabilities for divorce translate into expression at some threshold (e.g. [12]). Such models also permit estimation of ‘total heritability’ of divorce, resulting from the combination of VA_♀,_ VA_♂,_ and COVA_♀♂_ [13], which represents the overall potential for micro-evolutionary responses to selection on the population-wide distribution of liabilities. Such analyses require phenotypic observations of divorce versus mate-fidelity, conditional on survival between consecutive breeding attempts, from diverse relatives [14]. We use 39 years of comprehensive observations of free-living song sparrows (M*elospiza melodia)* to estimate (*i*) population-level divorce rate, (*ii*) VA_♀,_ VA_♂_ and COVA_♀♂_ in divorce, and (*iii*) individual repeatability and the sex-specific and total heritability, thereby assessing the potential for ongoing evolution of divorce as a key reproductive strategy.

## MATERIAL AND METHODS

A resident population of individually colour-ringed song sparrows on Mandarte Island, Canada has been intensively studied since 1975. Each year, all individuals alive in late April (typical start of breeding [15]) are recorded in a comprehensive census to determine over-winter survival and pairing status (re-sighting probability >0.99 [16]), and all breeding attempts are closely monitored (ESMS1). Females and males form socially persistent pairings that cooperate to rear offspring (1–4 broods per year), but can form new pairings within and between breeding seasons following divorce or mate-death.

To identify cases of divorce versus mate-fidelity, we extracted each female’s lifetime sequence of breeding events (≥1 egg laid) where re-pairing could have occurred (i.e. she initiated a subsequent breeding event) and categorized these events as divorce, mate-fidelity or mate-death according to the pair-bond’s fate and whether her current mate was still alive during her subsequent event (ESMS1). Instances of mate-death were identified from daily field observations and April censuses, and excluded from our dataset.

We fitted two generalized linear mixed models to decompose total variance in liability for divorce. Model 1 estimated variances attributable to permanent effects of individual females (V_I♀_), males (V_I♂_) and of unique female-male social pairings (V_S_), and the year when a focal breeding event occurred (V_Y_). Model 2 additionally estimated V_A♀_,V_A♂_, and COV_A♀♂_ in liability for divorce, using comprehensive pedigree data to quantify relatedness (i.e. an ‘animal model’ [9]; ESMS2). COV_A♀♂_ is the covariance between additive genetic effects of alleles expressed in all females versus all males, not the genetic covariance between a female and her socially-paired mate (e.g. [11,15]; ESMS3). Our current aim was to partition the total naturally-occurring variation in liability for divorce and thereby appropriately estimate heritability, not to explain variation in the occurrence of divorce. Fixed effects were consequently restricted to a two-level factor that defined whether an observation of divorce versus mate-fidelity spanned breeding events separated by the non-breeding season (‘between-season’) versus consecutive events within the same season (‘within-season’, ESMS1) and separate regressions on female and male individual coefficients of inbreeding (*f*) (Model 2 only), thereby estimating inbreeding depression in liability for divorce and minimising any associated bias in estimated V_A_ [17].

Sex-specific repeatabilities in liability for divorce were estimated from Model 1 as:

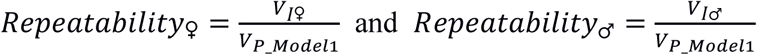

Sex-specific heritabilities (*h*^*2*^) were estimated from Model 2 as:

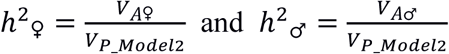

The ‘total heritability’ (*T*^*2*^) was calculated from Model 2 as:

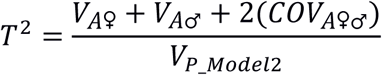

The total variance in liability (V_P_Model 1_, V_P_Model 2_) was calculated separately for each model (ESMS3). Models were fitted using Bayesian inference in `R`, using relatively uninformative priors (ESMS5). Posterior distributions of repeatabilities, heritabilities, and *T*^*2*^ were calculated from marginal posterior distributions of underlying variance components. We report liability-scale posterior modes, means and 95% credible intervals (95%CI) from 5000 posterior samples, and present prior distributions alongside posteriors to facilitate inference. Estimates and conclusions were robust to alternative model specifications (ESMS5).

## RESULTS

There were 1,419 breeding events where divorce could have occurred, involving 566 unique social pairings among 358 females and 341 males. Divorce occurred on 166 (11.7%) occasions (details in ESMS1).

Model 1 revealed that the largest component of variance in liability for divorce was V_I♀_, while V_I♂_ was comparatively small (table 1, ESMS4). Because V_S_ and V_Y_ were also small, female repeatability for divorce was moderate (∼16%; table 1). The lower 95%CI limit converged towards zero, but 98% of posterior samples exceeded 0.01, departing from the prior distribution (figure 1*a*), and indicating that female repeatability is most likely greater than zero. In contrast, male repeatability was smaller (table 1); only 76% of posterior samples exceeded 0.01 (figure 1*b*). Divorce was less likely to occur within a breeding season than between seasons (table 1).

**Table 1:** Marginal posterior modes, means and 95%CIs from models decomposing total variance in liability for divorce. V_I_ and V_A_ represent permanent individual and additive genetic variances for females (♀) and males (♂). COV_A♀♂_ is the cross-sex genetic covariance. V_S_ and V_Y_ are the social pair and year variances, respectively. Posterior statistics for sex-specific repeatabilities and heritabilities (*h*^*2*^), ‘total heritability’ (*T*^*2*^), fixed effects of within-season versus between-season (intercept) and regressions on individual female or male coefficient of inbreeding (*f*) are also shown.

|  | Model 1 |  | Model 2 |  |
| --- | --- | --- | --- | --- |
|  | mode, mean | 95%CI | mode, mean | 95%CI |
| **variance components** |  |  |  |  |
| $V_{I♀}$ | 0.24, 0.28 | $4 \times 10^{-6}$, 0.53 | 0.003, 0.22 | $6 \times 10^{-7}$, 0.48 |
| $V_{I♂}$ | 0.001, 0.09 | $5 \times 10^{-8}$, 0.26 | 0.002, 0.08 | $4 \times 10^{-8}$, 0.25 |
| $V_S$ | 0.001, 0.14 | $2 \times 10^{-9}$, 0.46 | 0.003, 0.15 | $4 \times 10^{-8}$, 0.48 |
| $V_Y$ | 0.001, 0.07 | $4 \times 10^{-7}$, 0.19 | 0.001, 0.07 | $8 \times 10^{-9}$, 0.19 |
| $V_{A♀}$ | | | 0.001, 0.07 | $5 \times 10^{-9}$, 0.25 |
| $V_{A♂}$ | | | 0.001, 0.08 | $4 \times 10^{-8}$, 0.21 |
| $COV_{A♀♂}$ | | | -0.0002, -0.003 | -0.08, 0.06 |
| **variance ratios** |  |  |  |  |
| repeatability$_{♀}$ | 0.16, 0.17 | $3 \times 10^{-6}$, 0.30 | | |
| repeatability$_{♂}$ | 0.001, 0.05 | $3 \times 10^{-8}$, 0.15 | | |
| $h^2_{♀}$ | | | 0.001, 0.04 | $2 \times 10^{-9}$, 0.14 |
| $h^2_{♂}$ | | | 0.001, 0.04 | $3 \times 10^{-8}$, 0.12 |
| $T^2$ | | | 0.02, 0.08 | $1 \times 10^{-4}$, 0.20 |
| **fixed effects** |  |  |  |  |
| intercept | -0.64, -0.66 | -0.89, -0.44 | -0.81, -0.90 | -1.28, -0.59 |
| within-season | -1.08, -1.10 | -1.34, -0.85 | -1.11, -1.12 | -1.37, -0.87 |
| $f_{♀}$ | | | 1.92, 1.98 | -0.66, 4.45 |
| $f_{♂}$ | | | 0.59, 0.94 | -1.72, 3.61 |

**Figure 1.**
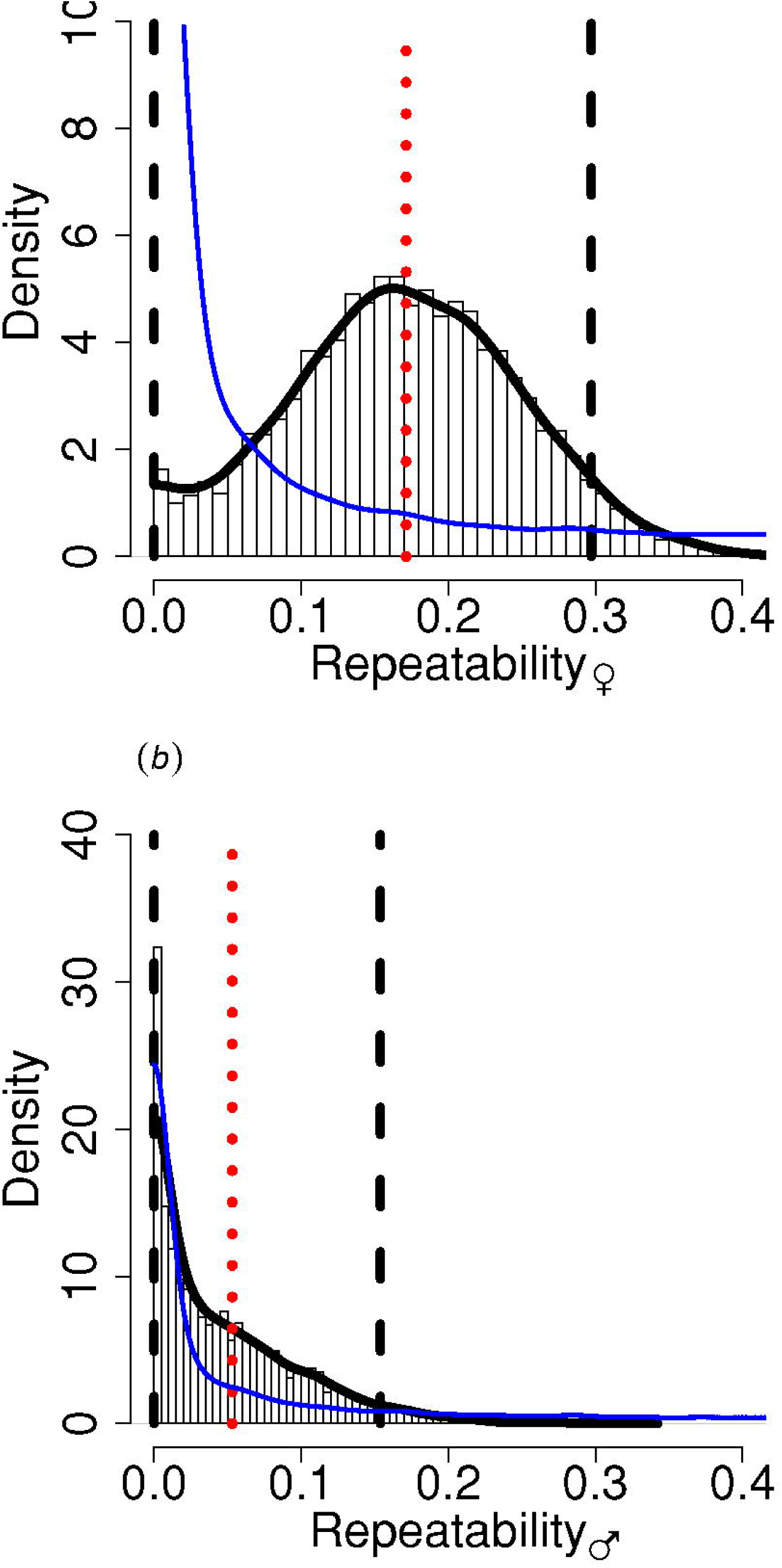
Marginal posterior samples (bars), density (solid black line), mean (red dotted line) and 95%CI limits (dashed lines) of sex-specific repeatabilities for divorce in (*a*) female and (*b*) male song sparrows. Blue lines illustrate prior distributions (ESMS5). Areas where the posterior density exceeds the prior density highlight parameter values that are likely supported by the data.

Model 2 showed that V_A♀_ and V_A♂_ in liability for divorce were both small, and COV_A♀♂_ was estimated as close to zero (table 1, ESMS4). Sex-specific heritabilities were therefore small for both females and males. However, posterior distributions departed from the priors, with higher density at higher values (e.g. at the posterior means, figure 2*a,b*) implying that heritabilities, and underlying V_A♀_ and V_A♂_, exceed zero. Indeed, V_A♀_, V_A♂_, and COV_A♀♂_ combined to generate a small but likely non-zero total heritability (*T*^*2*^) for divorce (table 1); 92% of posterior samples exceeded 0.01 (ESMS5). This again deviates from the prior distribution (figure 2*c*), and from the posterior distribution that would have resulted given zero V_A♀_, V_A♂_ and hence *T*^*2*^ (ESMS5). Liability for divorce tended to increase with increasing *f*, especially in females, but the 95%CIs overlapped zero (table 1).

**Figure 2.**
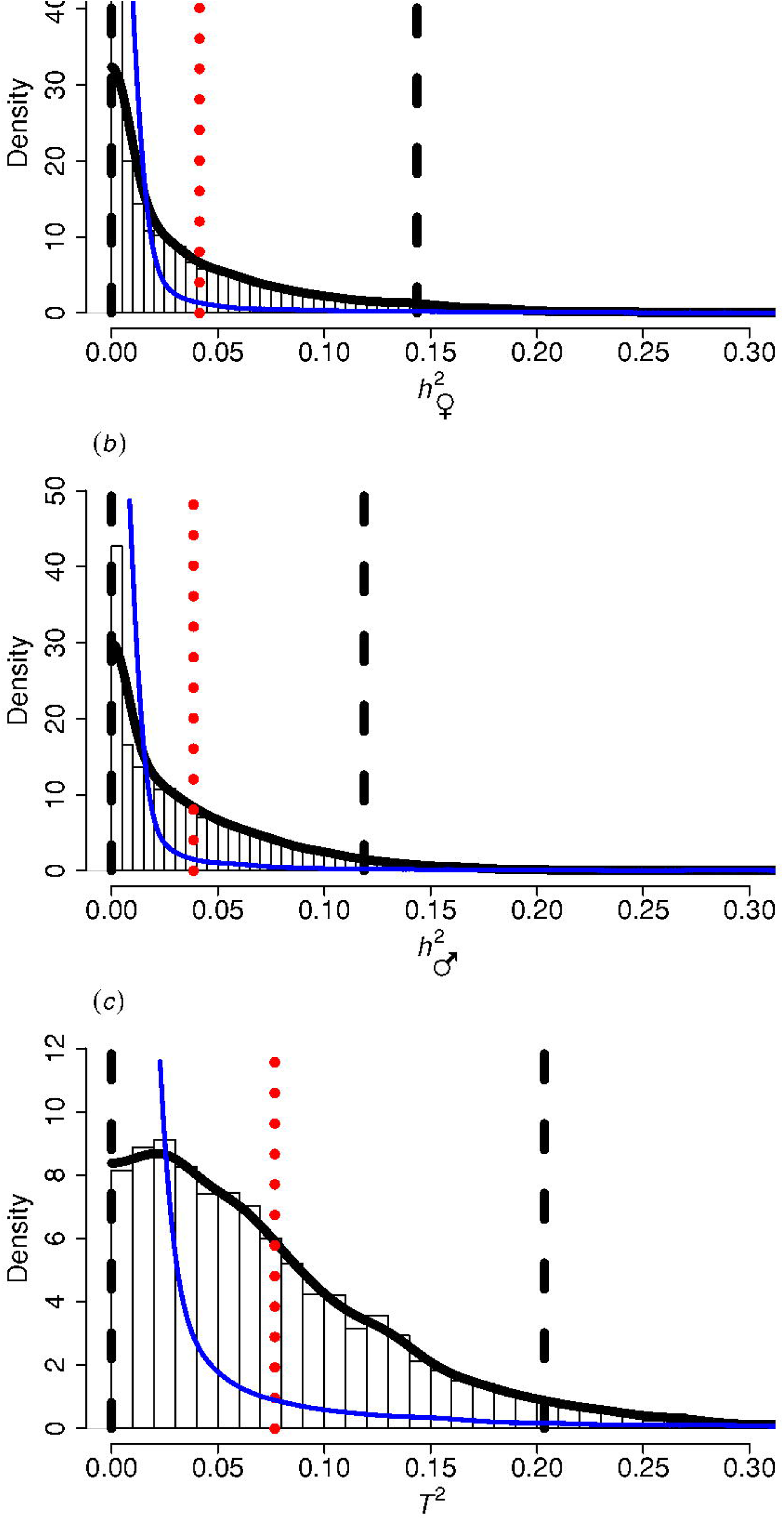
Marginal posterior distributions for (*a*) female and (*b*) male heritabilities (*h*^*2*^), and (*c*) the total heritability (*T*^*2*^) for divorce in song sparrows. See figure 1 for plot description.

## DISCUSSION

The ∼12% divorce rate observed in song sparrows is relatively low compared to other temperate-breeding passerine birds (∼20–50%; [14,18,19]). However, there was evidence of moderate V_I♀_ and hence female repeatability, but lower V_I♂_ and male repeatability, in liability for divorce (figure 1*a,b*, ESMS4). These estimates imply that sex-specific *h*^*2*^ is not *a priori* zero. However in practice, *h*^*2*^ was estimated to be small in both sexes.

Most previous quantitative genetic analyses of divorce come from human twin-studies, and show relatively high heritabilities with divorce defined as a sex-specific trait (e.g. 0.3–0.6 [20,21]). However, such estimates may be inflated by shared environmental and cultural effects [9], and often only consider whether individuals ever divorced over their lifetime. Our focus on sequential breeding events, considering among-individual variances across repeat observations, allows estimation of individual repeatability as well as pair and year variances, which encompass variances stemming from ecological and/or social environmental effects that could influence the occurrence of divorce. The only previous quantitative genetic analysis of divorce in a wild (non-human) population also estimated low female heritability in savannah sparrows (*Passerculus sandwichensis*) [14]. Because [14]’s estimate of male repeatability for divorce was not distinguishable from zero, heritability estimates were restricted to females and did not consider potential contributions of V_A♂_ or COV_A♀♂_. Our results suggest that both song sparrow sexes do contribute to the total additive genetic variance, and hence to the total heritability for divorce (*T*^*2*^). Indeed small, potentially undetectable, effects in each sex can combine to generate detectable total *T*^*2*^ [13]. Thus the overall apparent potential for micro-evolutionary responses to selection on divorce is greater when considering the interactive effects of the sexes jointly (i.e. considering divorce as an emergent trait) than when considering either sex alone.

Many studies have investigated the potential costs and benefits of divorce in wild populations, particularly in socially-monogamous birds (reviews: [2,3,5]). Divorce is generally considered to be adaptive, and increases an individual’s subsequent breeding success and hence fitness under certain conditions [2,3]. However, ongoing micro-evolutionary responses to such contemporary selection require V_A_. Our results indicate that such V_A_, and consequent potential for evolutionary response to selection, while probably non-zero, is smaller than is often implicitly assumed [2,4–6]. Further, relatively low divorce rates, such as observed in song sparrows, will intrinsically limit the intensity of selection [12]. Overall, rapid and marked micro-evolutionary changes in the frequency of divorce appear unlikely, even if divorce were beneficial for one or both members of a breeding pair. Consequently, there is also limited potential for genetic covariance between liability for divorce and other key reproductive strategy traits, such as female extra-pair reproduction (EPR), negating the suggestion that divorce and EPR both represent manifestations of an underlying ‘weak-pair’ syndrome [22] and limiting the potential for indirect selection.

## Ethics

All field procedures approved by the University of British Columbia Animal Care Committee ethical review (certificates UBCACC A12-0229 and A14-0336).

## Data accessibility

Data and code are available as electronic supporting information.

## Authors’ contributions

RRG and MEW designed the study and conducted analyses with input from JMR. All authors contributed to writing, agree to be held accountable for all aspects of the work, and approve the final version of the manuscript.

## Acknowledgments

We thank the Tsawout and Tseycum First Nation bands for access to Mandarte, Peter Arcese, Lukas Keller, Pirmin Nietlisbach, and the University of Aberdeen Maxwell High Performance Computing cluster.

