## Supplementary Materials for "Individual repeatability and heritability of divorce in a wild population"

**Electronic Supplemental Materials for:**

### Electronic Supplemental Material S1 – Descriptive details of re-pairing dataset

#### *S1.1 Classifying re-pairing due to divorce and mate-death*

Divorce rates are often highest in relatively short-lived species where re-pairing due to mate-death is also common [1]. However, because most study systems lack comprehensive observations of individual survival and/or records of the lifetime reproductive attempts, many investigations of divorce are severely limited in their ability to accurately ascribe instances of re-pairing due to divorce versus mate-death [2,3]. In contrast, the small size of Mandarte Island (~6ha) and intensity of field monitoring (below), mean that the causes of re-pairing among song sparrows can be accurately ascribed, presenting a valuable opportunity to dissect the quantitative genetic basis of divorce independent of mate-death.

In the Mandarte Island study population, males and females can begin breeding in their first adult season (aged one year) and rear 1–4 broods per year over a median lifespan of approximately 2 years [4]. Following a complete annual census in late April designed to document over-winter survival and identify new immigrant breeders, all individuals on Mandarte are closely monitored through the end of breeding in late July-early August [4]. All offspring surviving ~6 days post-hatch and immigrants ($0.92^{\text{-year}}$ on average) are colour-ringed. During monitoring, the territories of each breeding pair, and of unpaired males (typically ~10–40% of males annually), are visited every 3–5 days to record nesting and territorial behaviour. Any instances of territorial take-overs or within-season re-pairing are thus observed soon after they occur [5].

Each breeding event (social pairing where ≥1 egg was laid) constitutes an observation after which a social pair could potentially express divorce or mate-fidelity (conditional on both individuals surviving to a subsequent breeding event). To extract these observations, we tabulated each female's lifetime sequence of breeding events, except her last (where expression of mate-fidelity or re-pairing is not possible), and recorded whether the identity of her social mate changed between the focal and subsequent event. Instances where social male identity did not change were classified as 'mate-fidelity'. For instances where the female's social mate did change, we determined whether her mate from the focal event was observed in the population (breeding or otherwise) later that season during daily monitoring and/or survived to the following April census (i.e. within-season vs between-season re-pairing). In instances where the male survived to the next breeding event of his former mate, re-pairing was classified as 'divorce', whereas if the male did not survive re-pairing was classified as 'mate-death'. Breeding events ending in mate-death were excluded from the 'divorce dataset' (See ESMS1.2), but subsequent breeding events by the focal female which could have ended in divorce or mate-fidelity were retained. All observations in the dataset therefore describe whether a pair divorced or not after a given breeding event, and thus the focal trait describes a change in state between two consecutive breeding events. We attributed observations of divorce or mate-fidelity to characteristics of the initial breeding event of each social pair. Thus, independent variables associated with the trait (e.g. within- versus between-season, and observation year) are attributed to the initial event. The same phenotypic dataset would have been obtained had we instead focused on sequences of male rather than female breeding events [6,7].

Due to reduced field effort in 1980, information on social pairing throughout the breeding season and thus within-season divorce or mate-death was not available. However, information on the survival of all individuals alive at the beginning of the 1980 season was still collected via the April census. Consequently, two breeding events at the end of 1979 (i.e. each female's last breeding event for that year) could be accurately classified as 'between-season mate-death', whereas all other breeding events that could have ended in re-pairing at the end of 1979 (n = 5) were excluded due to uncertainty in whether females maintained mate-fidelity or re-paired in 1980.

#### *S1.2 Descriptive statistics of re-pairing dataset*

Of the 1,595 total breeding events which could have resulted in re-pairing from 1979–2015, there were 176 instances of re-pairing due to mate-death (11%). Of these, 120 mate-deaths occurred 'between-seasons', while 56 mate-deaths occurred 'within-season'. Of the 166 instances of divorce (see *Results*), 91 occurred 'between-seasons', while 75 occurred 'within-season'.

The 358 females included in our 'divorce dataset' (i.e. 1,419 breeding events where divorce could have occurred, *Results*) formed a mean of 1.58 (±0.90 SD, range 1–6) unique social pairings, and 220 females formed only one social pair. The 341 males formed a mean of 1.66 (±0.97 SD, range 1–7) unique social pairings, and 204 males formed only one social pair. Further, there were 72 exclusive pairings where a females only ever paired with a single male that did not pair with any other female. These characteristics of the data were sufficient to separate female and male individual effects on divorce in Models 1 and 2, as well as to separate social pair identity effects from the individual effects of each female and male (see ESMS5.3). Over the 36 observation years included in our study (1979–2015; excluding 1980, above), there was a mean of 39.4 (±21.9 SD, range 1–81) breeding events per year that could have ended in divorce. Of these, there was a mean of 4.6 (±3.1 SD, range 0–11) divorces per year.

**Electronic Supplemental Material S2 – Summary of pedigree construction**

We compiled a complete social pedigree, wherein detailed field observations from 1975–2015 were used to assign all ringed chicks to their social parents. All chicks ringed since 1993 and their potential parents, plus an additional sample of chicks ringed between 1986–1992, were genotyped at 160 polymorphic microsatellite loci, allowing genetic paternity assignment with extremely high confidence [8]. We used all available genetic parentage data to correct the social pedigree for ~28% extra-pair paternity so far as feasible (details of pedigree reconstruction [6]). The resulting pedigree for 1975–2015 was pruned to all phenotyped individuals and their known ancestors, and then used to construct the inverse of the numerator relatedness matrix and to calculate individual coefficients of inbreeding ($f$) (e.g. [9–11]). The pruned pedigree comprised 959 individuals. Across the phenotyped individuals, mean female $f$ was 0.054±0.053 SD, and mean male $f$ was 0.054±0.056 SD.

### Electronic Supplemental Material S3 – Model details and implementation

*S3.1 Model details*

The occurrence of divorce was modelled as a threshold trait using generalized linear mixed models, where each observation (i.e. mate-fidelity or divorce after a focal breeding event) is assigned a value for the normally distributed liability of divorce [12–14]. Such threshold models are a well-established method for treating dichotomous traits in quantitative genetics [12]. Conceptually, they consider an underlying continuously distributed (Gaussian) variable – the 'liability' – that translates into expression of a discrete phenotype at some threshold value. Such models ensure that the key quantitative genetic assumption of multivariate normality of additive genetic effects is fulfilled irrespective of the observed frequency of occurrence of the focal phenotype(s).

In our model, liability values below the threshold result in expression of mate-fidelity, whereas values above the threshold result in divorce. Liability is a joint trait of each focal social pair and is modelled by a linear function of factors unique to each female, male and social pair [8–10,14]. We fitted two nested models that decomposed the vector containing each pair's liability for divorce across observations ($\mathbf{l}$):

$$\mathbf{l} = \mathbf{X\beta} + \mathbf{Z_i i} + \mathbf{Z_s s} + \mathbf{Z_y y} \qquad \text{(Model 1)}$$

$$\mathbf{l} = \mathbf{X\beta} + (\mathbf{Z_a a} + \mathbf{Z_i i}) + \mathbf{Z_s s} + \mathbf{Z_y y} \qquad \text{(Model 2)}$$

into vectors of fixed effects ($\boldsymbol{\beta}$) and random permanent individual ($\mathbf{i}=[\mathbf{i}_{♀}', \mathbf{i}_{♂}']$), social pair identity ($\mathbf{s}$), year of initial breeding event ($\mathbf{y}$), and additive genetic ($\mathbf{a}=[\mathbf{a}_{♀}', \mathbf{a}_{♂}']$) effects. Parentheses in Model 2 are not mathematically necessary, but visually group sources of repeatable individual effects. Random effects $\mathbf{s}$ and $\mathbf{y}$ follow univariate normal distributions, defined by means of zero and variances to be estimated. Residual effects are not explicitly included in the liability of the threshold model (eqn. 4 of [14]). Female and male permanent individual and additive genetic effects are assumed to follow bivariate normal distributions, defined by means of zero and covariance matrices:

$$\boldsymbol{G_I} = \begin{bmatrix} V_{I♀} & 0 \\ 0 & V_{I♂} \end{bmatrix} \qquad \boldsymbol{G_A} = \begin{bmatrix} V_{A♀} & COV_{A♀♂} \\ COV_{A♀♂} & V_{A♂} \end{bmatrix}$$

where $V_{I♀}$ and $V_{I♂}$ are the variances among female and male permanent individual effects, $V_{A♀}$ and $V_{A♂}$ are the variances among female and male additive genetic effects, respectively, and $COV_{A♀♂}$ is the cross-sex covariance in additive genetic effects. Because related females and males, each socially paired to other individuals, share some alleles, $COV_{A♀♂}$ represents the covariance in additive genetic effects among opposite sex relatives, weighted by the probability of sharing alleles identical by descent. Note that $COV_{A♀♂}$ is not the covariance in additive genetic effects between a female and her socially paired mate [9,11]. However, no such cross-sex covariance among permanent individual effects (**i**) is defined. To estimate $\mathbf{G_A}$, individual female and male identities in Model 2 were each linked to the inverse of the numerator relatedness matrix [12,16] constructed from the pedigree (ESMS2, [8–11]). The threshold on the liability scale is set to zero by using a probit link function and fixing the residual variance to one [14]. Accordingly, this model simultaneously estimates female and male additive genetic effects on liability for divorce, and estimates the cross-sex covariance. It consequently does not *a priori* define divorce as primarily a trait of one sex or the other, or hence specify the sex-specific genetic effects as 'direct' or 'indirect' (*sensu* [17]). These estimated variance components partition the total variance in liability for divorce. They thereby encompass effects of ecology and the social environment on that liability such as could stem, for example, from variation in the presence of other potential mates.

Because our aim was to estimate repeatability and heritability for the observed variation in liability for divorce, not to explain population-wide variation in divorce rate or partition variance in divorce conditioned on all possible environmental influences, we fitted minimal fixed effects. Specifically, we modelled between-season versus within-season effects and regressions on individual *f* (see *Methods*), but did not explicitly model effects of other variables, for example including pair relatedness (for which see [7]). To minimize under-estimation of *f*, phenotypic observations were restricted to individuals with known grandparents, hence truncated to 1979–2015, and excluded immigrants.

#### *S3.2 Model implementation*

Models were fitted using Bayesian inference to obtain 5,000 samples of the posterior distribution of parameters using the R [18] packages `MCMCglmm` [13] and `nadiv` [19], following a burn-in of 10,000 iterations and implementing a thinning interval of 5,000 iterations for a total of 25,010,000 model iterations. We used diffuse normal prior distributions (mean=0 and variance=$10^{10}$) for all fixed effects and parameter expanded priors for variance components, which gave relatively uninformative priors with scaled non-central *F*-distributions of numerator and denominator degrees of freedom equal to one [20] and scale parameter of 1,000.

Since liability for divorce was modelled as an emergent ('joint') trait stemming from the individual (Models 1 and 2) and additive genetic (Model 2) effects of both females and males, as well as the additive genetic covariance (Model 2), total phenotypic variance ($V_P$) for divorce in each model, conditioned on the fitted fixed effects, is approximated on the liability scale as:

$$V_{P\_\text{Model 1}} = V_{I\text{♀}} + V_{I\text{♂}} + V_S + V_Y + 1 \qquad \text{(Model 1)}$$

$$V_{P\_\text{Model 2}} = V_{A\text{♀}} + V_{A\text{♂}} + 2(COV_{A\text{♀♂}} \times \bar{r}) + V_{I\text{♀}} + V_{I\text{♂}} + V_S + V_Y + 1 \qquad \text{(Model 2)}$$

Here, $\bar{r}$ is the mean female-male relatedness across all observed breeding pairs, where relatedness $r$ for each observed pairing is calculated as twice the coefficient of kinship between paired individuals, as obtained from the numerator relatedness matrix constructed from the pedigree [21–23]. Across the 566 pairings that contributed phenotypic data, $\bar{r}$ = 0.146. The addition of a one on the right hand side of each equation for $V_P$ is the link scale variance for a probit link function and additive overdispersion model. Unlike in a binomial model with logit link, no residual variance is included in the calculation of $V_P$ for a threshold model with probit link, because the residual effect is incorporated as part of the threshold parameter (see the description of Model 2 above and eqn. 4 of [14]).

**Electronic Supplemental Material S4 – Full posterior variance component distributions**

**Figure S1.** Marginal posterior samples (bars), density (solid black line), mean (red dotted line), and 95%CI limits (dashed lines) of variance component estimates from Model 1, consisting of (*a*) female permanent individual variance ($V_{I♀}$), (*b*) male permanent individual variance ($V_{I♂}$), (*c*) year variance ($V_Y$), and (d), variance due to unique social pairings ($V_S$). Blue lines illustrate the marginal prior density (see ESMS5.1).

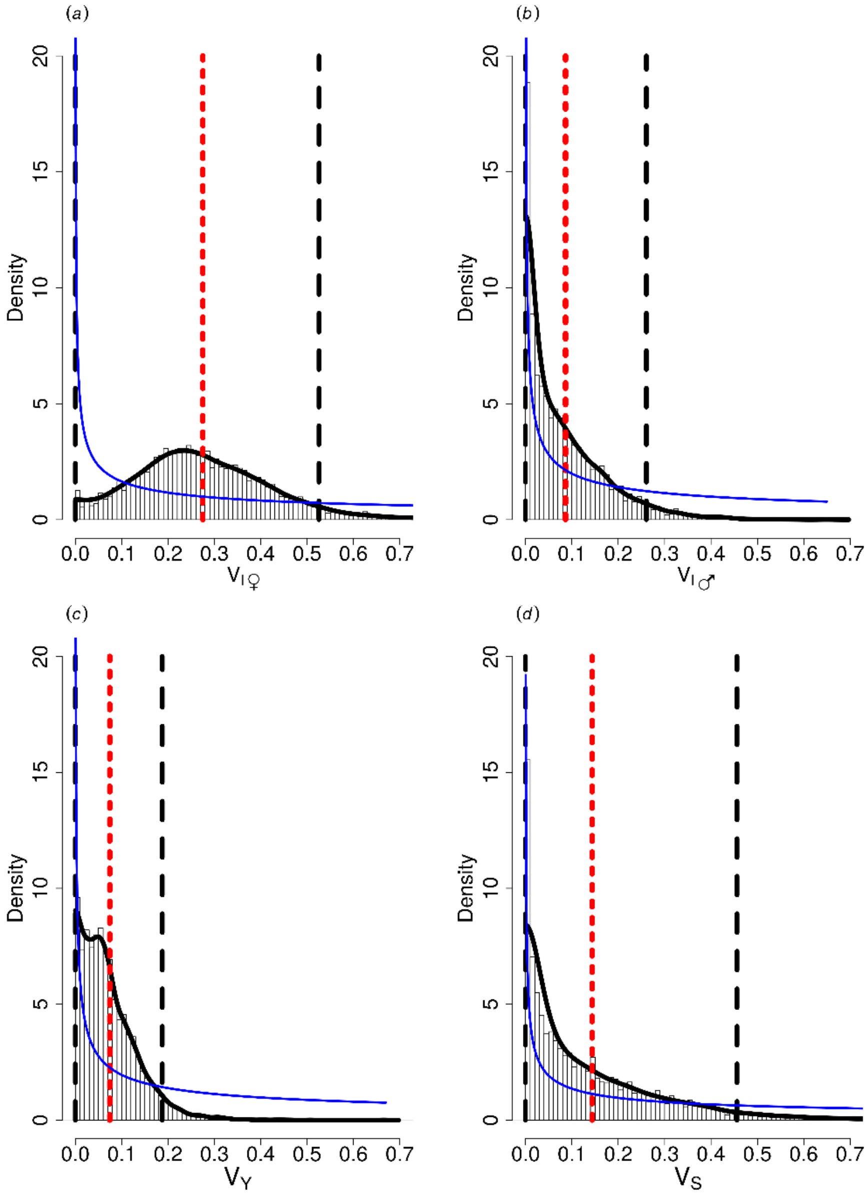

**Figure S2.** Marginal posterior distributions of variance component estimates from Model 2 (see figure S3 for additive genetic components), consisting of (*a*) female permanent individual variance ($V_{I♀}$), (*b*) male permanent individual variance ($V_{I♂}$), (*c*) year variance ($V_Y$), and (d), variance due to unique social pairings ($V_S$). See figure S1 for plot description.

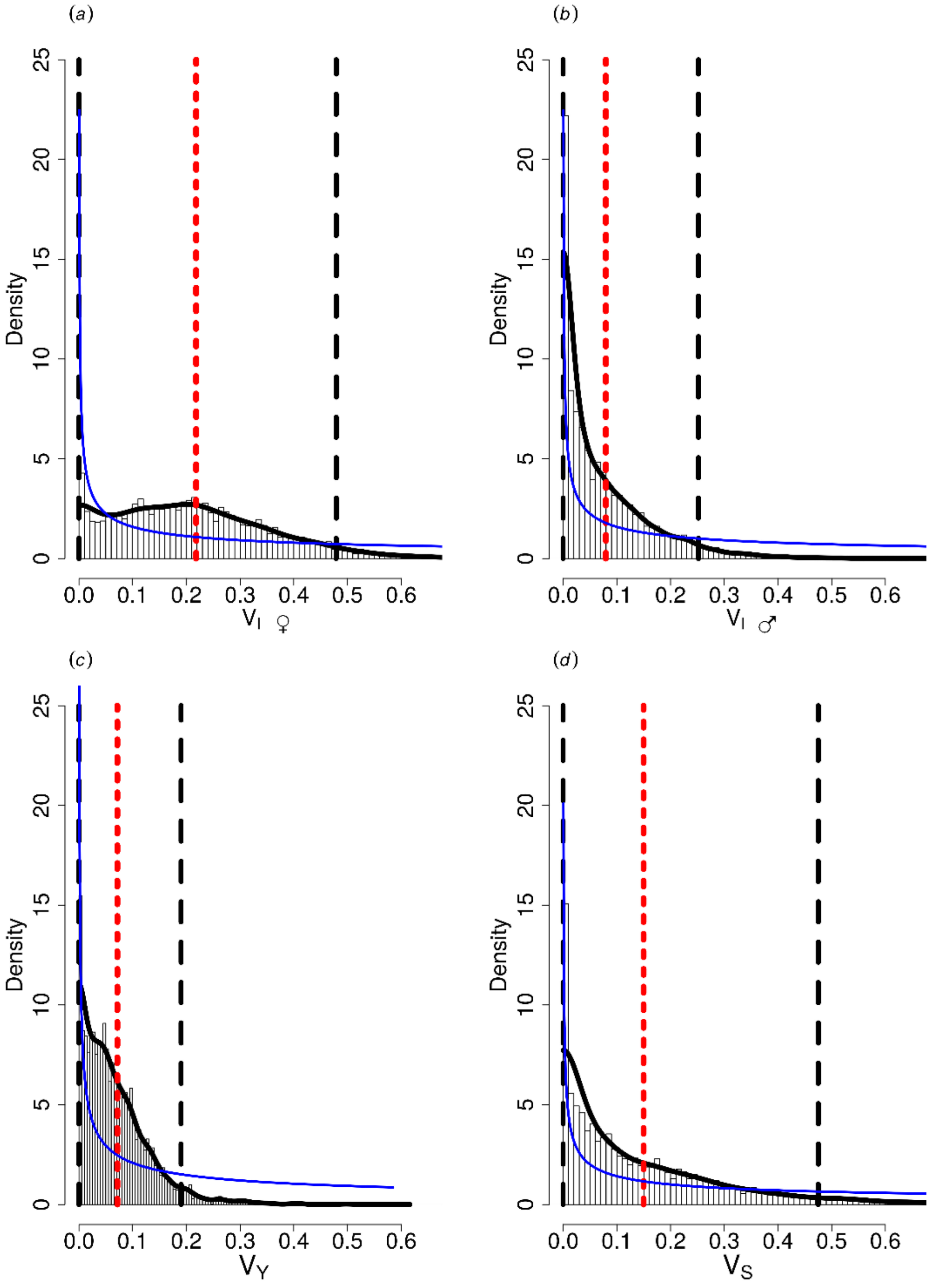

**Figure S3.** Marginal posterior distributions of all additive genetic component estimates from Model 2, consisting of (*a*) female additive genetic variance ($V_{A♀}$), (*b*) male additive genetic variance ($V_{A♂}$), (*c*) the cross-sex additive genetic covariance ($COV_{A♀♂}$), and *(d)* total additive genetic variance ($V_{A\text{-Total}}$). See figure S1 for plot description and ESMS5.3 for notes about the shape of $V_{A\text{-Total}}$. Note different scale of x and y-axes for (*c*).

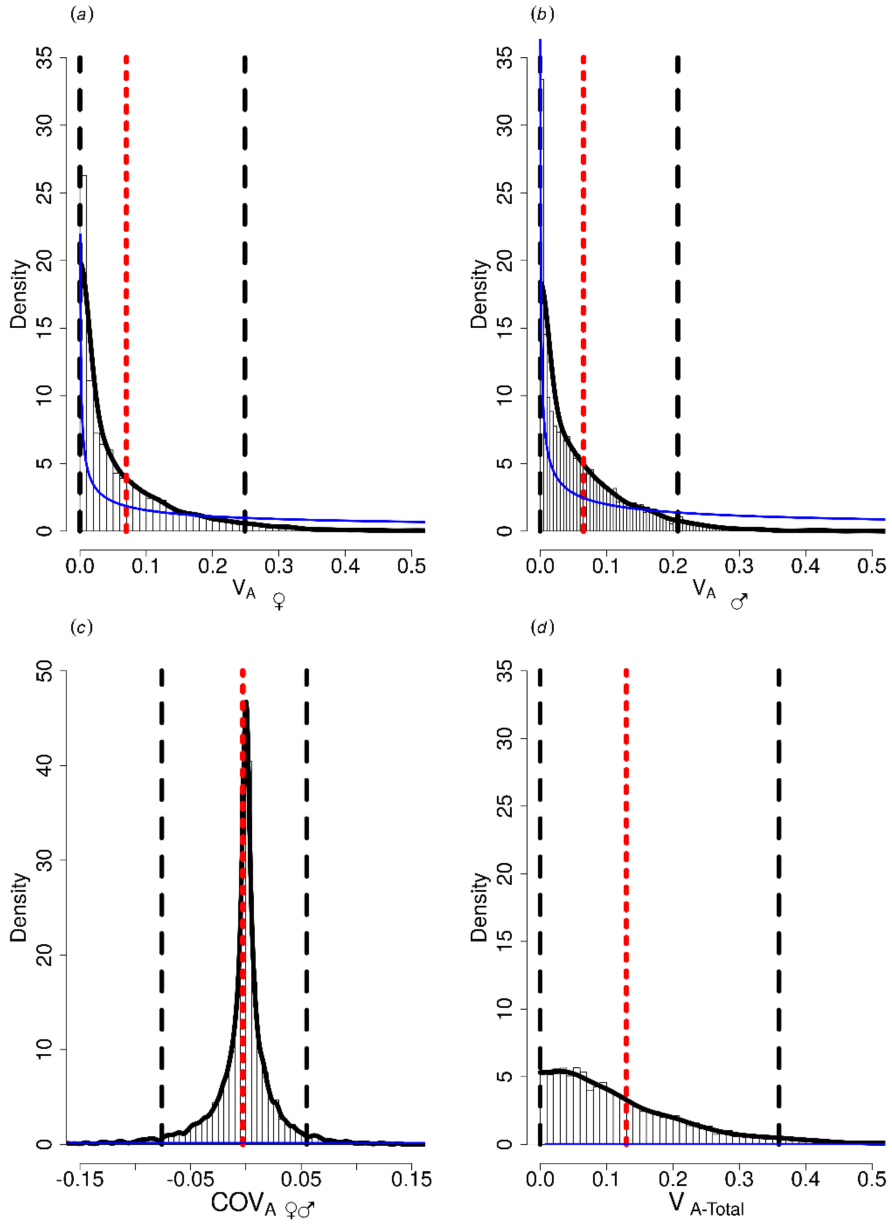

**Electronic Supplemental Material S5 - Alternative model results**

*S5.1 Prior influence on posterior distributions*

Since we had no strong *a priori* prediction regarding the magnitudes of effects on liability for divorce, we specified relatively uninformative priors. To visually inspect the influence of prior specifications on posteriors we plotted an approximate prior density on top of each plot of posterior samples. For all independent variance components, prior densities were derived by evaluating the probability density function for the *F*-distribution (section 3.6 of [13]) at each of the posterior histogram mid-points. These densities were then scaled to approximate a total area of one under the prior density curve.

For all covariances and associated variances, approximate prior densities were obtained from 10,000 random draws from the prior distribution. For the repeatabilities and heritabilities, approximate prior distributions were obtained by performing the same calculations as for the posterior with each sample of the simulated prior. A constant of one was added to the denominators for repeatability and heritability to account for the constant probit link variance that was introduced into the calculations of these values on the liability scale (see ESMS3.2). Kernel density estimation was then applied to the samples from the prior. Plot areas where posterior samples have a different shape from the prior distribution (e.g. more/less density in a tail, or less density under the mode) indicate that the posterior distribution is likely influenced by the data rather than solely by the prior.

*S5.2 Pre-1993 pedigree error due to extra-pair paternity*

While extra-pair paternity before 1993 presumably introduces error into the 1975–1992 pedigree, approximately 90% of all pedigree links are likely to be correct [6,10,11]. Such error likely causes little bias in estimates of $V_A$ [24], even if paternity error is non-random [25]. To verify this expectation, we additionally fitted Model 2 to a data subset comprising phenotypic and pedigree data starting from individuals hatched from 1993 onwards to ensure all phenotyped individuals had genetically verified parents. The restricted phenotypic data comprised 661 breeding events where divorce could have occurred during 1994–2015, containing 305 unique social pairings involving 192 females and 189 males. Divorce occurred on 92 (13.9%) occasions.

We altered the pedigree by removing all individuals with parents that were not genetically assigned, and pruned this pedigree to the restricted set of phenotyped individuals. Thus, individuals with genetically un-assigned parents, but that were the genetically assigned parents of a subsequent generation, became the founder individuals. Note that because total variance on the liability scale depends on the observed phenotypic mean (i.e., divorce rate), results are not quantitatively comparable between the full and restricted datasets. However, the pattern of variance partitioning among the different components is similar between the two models (table S1). This supports the view that pre-1993 pedigree error does not substantively bias estimates of additive genetic (co)variances. Model 1, and resulting estimates of repeatabilities, do not rely on a pedigree and so will not be affected by pedigree errors.

**Table S1:** Marginal posterior modes, means and 95%CIs of variance components from models decomposing the liability for divorce in the full dataset (main text Model 2) or data since 1993. $V_I$ and $V_A$ represent permanent individual and additive genetic variances for females (♀) and males (♂), and $COV_{A♀♂}$ is the cross-sex genetic covariance. $V_S$ and $V_Y$ are the social pair and year variances, respectively.

| | Main text Model 2 | | Post-1993 data and pedigree Model 2 | |
|---|---|---|---|---|
| | **mode, mean** | **95%CI** | **mode, mean** | **95%CI** |
| **variance components** | | | | |
| $V_{I♀}$ | 0.003, 0.22 | $7×10^{-7}$, 0.48 | 0.003, 0.29 | $2×10^{-8}$, 0.86 |
| $V_{I♂}$ | 0.002, 0.08 | $4×10^{-8}$, 0.25 | 0.002, 0.18 | $1×10^{-8}$, 0.61 |
| $V_S$ | 0.003, 0.15 | $4×10^{-8}$, 0.48 | 0.004, 0.42 | $2×10^{-7}$, 1.28 |
| $V_Y$ | 0.001, 0.07 | $8×10^{-9}$, 0.19 | 0.05, 0.18 | $1×10^{-7}$, 0.49 |
| $V_{A♀}$ | 0.001, 0.07 | $5×10^{-9}$, 0.25 | 0.001, 0.10 | $5×10^{-9}$, 0.38 |
| $V_{A♂}$ | 0.001, 0.08 | $4×10^{-8}$, 0.21 | 0.003, 0.16 | $1×10^{-8}$, 0.54 |
| $COV_{A♀♂}$ | -0.0002, -0.003 | -0.08, 0.06 | -0.0007, -0.006 | -0.14, 0.11 |

*S5.3 Alternative versions of Model 2: separation of sex-specific individual*

The total number of unique social pairings, pairings per individual, number of exclusive pairings (ESMS1.2), and previous analyses of song sparrow data [9–11], support the ability of our dataset to produce unbiased estimates of female and male $V_I$, $V_A$, and $COV_A$s in shared or emergent trait models.

In the current manuscript, the sampling correlation between $V_{A♀}$ and $V_{A♂}$ from Model 2 is 0.04, indicating that estimates of these two *variance components* are essentially independent of one another (the sampling covariance is a relationship between different quantities than $COV_{A♀♂}$; the latter measures the covariance between random additive genetic *effects* as expressed in a female versus a male). Because variances are bounded to be greater than zero, the small sampling correlation between $V_{A♀}$ and $V_{A♂}$ explains why the shape of the $T^2$ posterior differs noticeably from the shape of each of the $V_{A♀}$ and $V_{A♂}$ posterior distributions (figure 2, figure S3). In this circumstance, when $V_{A♀}$ is almost zero the sampling correlation implies that $V_{A♂}$ is likely to be relatively large, and vice versa, because $V_{A♂}$ is bounded at zero and so can only be uncorrelated with a small value of $V_{A♀}$ if $V_{A♂}$ is large. As $T^2$ is a function of the sum of $V_{A♀}$ and $V_{A♂}$, posterior samples where both $V_{A♀}$ and $V_{A♂}$ are close to zero will be infrequent. This explains why the high density at zero for $V_{A♀}$ and $V_{A♂}$ does not also occur for $T^2$ and why calculation of $T^2$ using posterior modes/means (i.e., summary statistics of the marginal posterior distributions) does not equate to the posterior mode/mean of the calculated $T^2$ for each sample of the posterior distributions.

The sampling correlation between $V_{I♀}$ and $V_{I♂}$ from Model 2 is -0.02 (Model 1 correlation: 0.0004), also indicating that estimates of these two variance components are independent from one another. Consequently, re-running Model 2 with divorce as either a female or male trait (i.e., removing $V_A$ and $V_I$ of the opposite sex mate in each pair) produced quantitatively similar posterior modes and means of the variance components (table S2).

Similarly, the sampling correlations between individual variances within each sex (i.e., $V_A$ and $V_I$) is -0.23 for females and -0.14 for males. Low correlations such as these indicate that reported variance component estimates for sex-specific effects on the liability for divorce are not confounded with each other, and hence not biased by other components in the model.

**Table S2:** Marginal posterior modes, means and 95%CIs of variance components from models decomposing the liability for divorce as either a female or male trait. $V_I$ and $V_A$ represent permanent individual and additive genetic variances for females (♀) and males (♂), and $COV_{A♀♂}$ is the cross-sex genetic covariance. $V_S$ and $V_Y$ are the social pair and year variances, respectively.

| | **Model 1** | | | | **Model 2** | | | |
|---|---|---|---|---|---|---|---|---|
| | **Female** | | **Male** | | **Female** | | **Male** | |
| | **mode, mean** | **95%CI** | **mode, mean** | **95%CI** | **mode, mean** | **95%CI** | **mode, mean** | **95%CI** |
| **variance components** | | | | | | | | |
| $V_{I♀}$ | 0.27, 0.28 | $5\times10^{-6}$, 0.51 | | | 0.002, 0.22 | $1\times10^{-10}$, 0.47 | | |
| $V_{I♂}$ | | | 0.001, 0.11 | $9\times10^{-7}$, 0.30 | | | 0.002, 0.10 | $2\times10^{-6}$, 0.30 |
| $V_S$ | 0.003, 0.17 | $1\times10^{-9}$, 0.50 | 0.38, 0.37 | $4\times10^{-7}$, 0.74 | 0.004, 0.18 | $4\times10^{-9}$, 0.51 | 0.21, 0.36 | $1\times10^{-5}$, 0.78 |
| $V_Y$ | 0.0008, 0.07 | $1\times10^{-7}$, 0.19 | 0.05, 0.09 | $2\times10^{-6}$, 0.22 | 0.001, 0.06 | $2\times10^{-7}$, 0.17 | 0.06, 0.09 | $4\times10^{-8}$, 0.23 |
| $V_{A♀}$ | | | | | 0.001, 0.06 | $3\times10^{-9}$, 0.22 | | |
| $V_{A♂}$ | | | | | | | 0.001, 0.08 | $8\times10^{-9}$, 0.24 |

Similarly, we demonstrate that estimates of variance among individual repeatable effects ($V_I$) are not confounded with variance among pair-level repeatable effects ($V_S$) in Model 1. The sampling correlation between $V_S$ and $V_{I♀}$ is -0.22 and between $V_S$ and $V_{I♂}$ is -0.11. Results from a version of Model 1 without the social pair term ($V_S$) show no appreciable change in either $V_{I♀}$ or $V_{I♂}$ (table S3). In this model, the marginal posterior distribution of $V_{I♀}$ no longer converges towards zero, suggesting the slightly higher sampling correlation of $V_{I♀}$ with $V_S$, compared to $V_{I♂}$ and $V_S$, has a small effect on the precision of $V_{I♀}$. However, the posterior modes and means remained similar, implying that our estimates of female and male permanent individual effects, and hence repeatability, are robust.

**Table S3:** Marginal posterior modes, means and 95%CIs of variance components from models decomposing the liability for divorce with the full Model 1 or from a model with social pair variance ($V_S$). $V_I$ represents permanent individual variances for females (♀) and males (♂) and $V_Y$ is the year variance.

| | **Model 1** | | **Model 1 Without $V_S$** | |
|---|---|---|---|---|
| | **mode, mean** | **95%CI** | **mode, mean** | **95%CI** |
| **variance components** | | | | |
| $V_{I♀}$ | 0.24, 0.28 | $4\times10^{-6}$, 0.53 | 0.25, 0.30 | 0.065, 0.56 |
| $V_{I♂}$ | 0.001, 0.09 | $5\times10^{-8}$, 0.26 | 0.001, 0.10 | $2.23\times10^{-8}$, 0.29 |
| $V_S$ | 0.001, 0.14 | $2\times10^{-9}$, 0.46 | | |
| $V_Y$ | 0.001, 0.07 | $4\times10^{-7}$, 0.19 | 0.001, 0.06 | $7\times10^{-11}$, 0.16 |

*S5.4 Null Simulations of Model 2*

To further support our inference from Model 2 that liability for divorce exhibits small but likely non-zero sex-specific and total heritability, we conducted a Monte Carlo simulation to estimate key parameters from data where $V_{A♀}$ and $V_{A♂}$ in liability for divorce are zero.

For each of 50 replicates, we randomly sampled the observed incidences of divorce versus mate-fidelity without replacement, and replaced each observed value in our dataset with the randomised values. We thereby retained the observed divorce rate, and did not change the total variance to be partitioned.

We then fitted the Model 2 structure to each of the 50 resampled datasets exactly as for the real dataset, retaining the same random and fixed effects structures. The posterior mean sex-specific additive genetic variances and heritabilities, and the total heritability ($T^2$), were computed for each replicate, and the grand mean and standard deviation of these posterior means was calculated across the 50 replicates.

Across the simulated null datasets, the grand mean posterior mean sex-specific $V_A$ and $h^2$, and the total heritability ($T^2$), in liability for divorce were all smaller than the posterior mean values estimated given the real data (Table S4). The difference frequently exceeded two standard deviations, providing strong evidence that the estimates from the real data differ from those generated by the null simulations (even though, as expected given the intrinsically zero-bounded and right-skewed posterior distributions, the null posterior means were not zero). Consequently, we conclude that the posterior means estimated from the real data indicate that the sex-specific $V_A$ and $h^2$, and $T^2$, in liability for divorce are small but most likely greater than zero.

**Table S4**: Marginal posterior means from Model 2 fitted to the real data and the mean (and one standard deviation, SD) of the posterior means across 50 null simulations in which the occurrence of divorce was randomly sampled. $V_A$ represents additive genetic variances for females (♀) and males (♂). $COV_{A♀♂}$ is the cross-sex genetic covariance. Results for sex-specific heritabilities ($h^2$) and 'total heritability' ($T^2$) are also shown.

| | **Real data mean** | **'Null Simulation' mean (SD)** |
|---|---|---|
| **variance components** | | |
| $V_{A}$♀ | 0.07 | 0.023 (0.013) |
| $V_{A}$♂ | 0.08 | 0.020 (0.009) |
| $COV_{A}$♀♂ | -0.003 | $4\times10^{-5}$ (0.001) |
| **variance ratios** | | |
| $h^2$♀ | 0.04 | 0.020 (0.010) |
| $h^2$♂ | 0.04 | 0.017 (0.007) |
| $T^2$ | 0.08 | 0.036 (0.013) |

(*Melospiza melodia*). *Am. Nat.* **187**, 736–752. (doi:10.1086/686198)

10. Germain RR, Wolak ME, Arcese P, Losdat S, Reid JM. 2016 Direct and indirect genetic and fine-scale location effects on breeding date in song sparrows. *J. Anim. Ecol.* **85**, 1613–1624. (doi:10.1111/1365-2656.12575)
11. Reid J, Arcese P, Keller L, Losdat S. 2014 Female and male genetic effects on offspring paternity: additive genetic (co)variances in female extra-pair reproduction. *Evolution.* **68**, 3357–2370.
12. Lynch M, Walsh B. 1998 *Genetics and Analysis of Quantitative Traits*. Sunderland, MA: Sinauer.
13. Hadfield J. 2010 MCMC methods for multi-response generalized linear mixed models: the MCMCglmm R package. *J. Stat. Softw.* **33**, 1–22.
14. Hadfield JD. 2015 Increasing the efficiency of MCMC for hierarchical phylogenetic models of categorical traits using reduced mixed models. *Methods Ecol. Evol.* **6**, 706–714. (doi:10.1111/2041-210X.12354)
15. McAdam AG, Garant D, Wilson AJ. 2014 The effects of others' genes: maternal and other indirect genetic efffects. In *Quantitative Genetics in the Wild* (eds A Charmantier, D Garant, LEB Kruuk), pp. 84–103. Oxford: Oxford University Press.
16. Hadfield JD, Wilson AJ, Garant D, Sheldon BC, Kruuk LEB. 2010 The misuse of BLUP in ecology and evolution. *Am. Nat.* **175**, 116–125.
17. Moore AJ, Brodie ED, Wolf JB. 1997 Interacting phenotypes and the evolutionary process: I. Direct and indirect genetic effects of social interactions. *Evolution.* **51**, 1352–1362.
18. R Development Core Team. 2015 R: A Language and Environment for Statistical Computing.
19. Wolak ME. 2012 nadiv : an R package to create relatedness matrices for estimating

non-additive genetic variances in animal models. *Methods Ecol. Evol.* **3**, 792–796. (doi:10.1111/j.2041-210X.2012.00213.x)
